# Quantifying the dynamics of hematopoiesis by *in vivo* IdU pulse-chase, mass cytometry and mathematical modeling

**DOI:** 10.1101/552489

**Authors:** Amir Erez, Ratnadeep Mukherjee, Grégoire Altan-Bonnet

**Author notes:** These authors contributed equally to this work (Amir Erez, Ratnadeep Mukherjee).

## Abstract

We present a new method to directly quantify the dynamics of differentiation of multiple cellular subsets in unperturbed mice. We combine a pulse-chase protocol of IdU injections with subsequent analysis by mass cytometry (CyTOF), and mathematical modeling of the IdU dynamics. Measurements by CyTOF allow for a wide range of cells to be analyzed at once, due to the availability of a large staining panel without the complication of fluorescence spill-over. These are also compatible with direct detection of integrated iodine signal, with minimal impact on immunophenotyping based on surface markers. Mathematical modeling beyond a binary classification of surface marker abundance allows for a continuum of cellular states as the cells transition from one state to another. Thus, we present a complete and robust method for directly quantifying differentiation at the systemic level, allowing for system-wide comparisons between different mouse strains and/or experimental conditions.

## Introduction

The origin, development and turnover of leukocytes is a longstanding yet fundamental question in immunology. Fifty years ago, Ralph van Furth and Zanvil Cohn (1) used a pulse-chase technique to monitor the development of monocytes and macrophages. By using ^3^*H* incorporation and meticulous microscopy, they shaped our understanding of innate immunity by pointing out that tissue macrophages most likely originated from blood monocytes, that themselves originated from promonocytes in the bone marrow. Much has changed since the 1960’s yet the study of hematopoiesis and leukocyte development continues to provide interesting questions and intriguing answers. Importantly, the field of differentiation and hematopoiesis suffers from a noticeable gap between in-vivo and in-vitro studies. Whereas many in-vitro and cell-transplantation assays revealed what hematopoietic stem cells (HSC) and other leukocytes are *capable* of differentiating to, these studies do not tell us what actually happens in a healthy adult. Indeed, since cell-transplantation experiments usually involve depleting the host populations (by eg., irradiation), the resulting “empty” mouse needs to rapidly reconstruct its entire immune system, operating in highly inflammatory conditions. It is thus unclear how to apply knowledge gained from such perturbed conditions on healthy organisms at homeostasis.

Several approaches currently exist to study hematopoiesis, each with its merits and limitations. Ablation of relevant populations following a knockout of a gene, eg., a transcription factor such as FoxP3 (2), typically serve as conclusive evidence to define master regulators of leukocyte differentiation. Such genetic approach to probe leukocyte development, can be deceptive, as was recently demonstrated in the context of thymic negative selection of self-reactive T-cells (3). Alternatively, much can be gained by studying unperturbed healthy adult mice. Computational approaches (4, 5, 6, 7) use an algorithm for lineage inference without requiring time-series data, by hypothesizing that there is a continuous relationship between maturing cell subsets. As hematopoiesis is both continuous and asynchronous, the full spectrum of cell types exists in a single bone-marrow sample from a healthy individual. Though powerful, such approaches are currently limited to particular differentiation scenarios and lack a direct observation of the dynamics. Indeed, most of these approaches map out differentiation along a pseudo-time dimension based on the proximities of cell states.

In contrast, others (8) have combined inducible in-situ labeling (the so-called barcoding) and mathematical modeling to reveal the nature of the HSC stem cell contribution to hematopoietic maintenance. Specific labeling of HSC cells in mice is achieved by expression of a modified Cre recombinase from the Tie2(Tek) locus. Subsequent articles (9) have dissected the benefits of such in-situ genetic tagging in comparison to traditional techniques. An important advantage for in-situ labeling is its non-perturbative nature and its flexibility in assigning labeling targets. Crucially, to interpret the resulting time-series it has been essential to use mathematical modeling. However, such genetic manipulation is not easy or quick, and can severely limit the applicability of the approach on different strains or species.

A giant leap in the field of single-cell RNA sequencing demonstrated that dynamic maps of differentiation could be inferred using a clever model of RNA maturation and splicing. Kharchenko and colleagues (10) introduced a powerful approach to reconstitute the whole dynamics of RNA transcription and maturation in entire tissues or organisms. In turn, they used their RNA dynamic map to infer the dynamic relationship between the cells themselves. This method (called *velocyto*) indeed delivers “real-time” measurements of cell differentiation, when all previous methods in scRNAseq were confined to inferring pseudo-times. Yet, the discrepancies between RNA and protein studies(11) are warranting the development of dynamic methods in the field of mass cytometry (CyTOF) to leverage the protein-resolution of cytometry with the labeling depth of CyTOF.

Proliferation specific tags such as BrdU (*5-Bromo-2-Deoxyuridine*), EdU (*5-Ethynyl-2-Deoxyuridine*) and IdU (*5-Iodo-2’-deoxyuridine*), jointly referred to as XdU in this publication, offer an alternative for protein-based cytometry (FACS or mass cytometry). XdU gets inserted in the genomic material of cells, when they are undergoing DNA replication. Accordingly, proliferating cells (and only them) get tagged with XdU, and, upon division after tagging, cells pass on half their XdU to their daughters. The tagged cells can also differentiate by changing their phenotype (by *e.g*., expressing different surface markers), in which case they carry the XdU content, undiluted, to their new phenotypic compartment. XdU can be administered in the drinking water (12), or injected intraperitoneally or intravenously, followed by flow-cytometry and fitting of XdU time-series experiments to a mathematical model. The resulting time-series give insight about proliferating populations and their progeny, *in vivo*, without perturbing the animal and without requiring the elaborate genetics used in the in-situ fate-mapping techniques outlined in the previous section. Moreover, because administration of the tag stops at some point, the time-series can observe both the labeling and the de-labeling phases, thereby providing more information than the equivalent time-series from in-situ tags.

Upon intraperitoneal injection in mice, XdU is made bio-available for less than an hour after which it gets metabolized. Cells that are undergoing proliferation get tagged practically instantaneously; upon proliferation, they dilute their XdU content; upon differentiation, they bequeath the XdU tag to their descendant. In the context of monocyte differentiation, such a flow cytometry (FCM) based method has been previously used (13). However, FCM is disadvantaged by lack of breadth of markers to simultaneously study different cell groups. Moreover, antibody staining for BrdU requires harsh permeabilization of the cells (for the anti-BrdU antibody to reach its nuclear target) and this can be detrimental to the detection of various epitopes and surface markers. EdU requires a simpler protocol, yet relies on “click” chemistry which is incompatible with certain common dyes. Instead, here, we propose to use a combination of IdU injection followed by analysis using mass cytometry (CyTOF), which circumvents these problems. IdU is immediately detected as elemental Iodine in the CyTOF without the requirement of any additional experimental step. The CyTOF allows for a wide staining panel, capturing several cell groups in detail and providing a systemic view of differentiation. Whereas IdU injection in mice has been used in microscopy imaging, for neogenesis (14), cell cycle analysis (15), and proliferation (16), its use in single-cell mass cytometry remains limited. IdU has been used *in vitro* on CyTOF measurements (17) to study cell cycle and to immunologically profile proliferating versus non-proliferating cells in the mouse bone marrow (18), however, it remains an untapped resource to study differentiation. Even so, studying hematopoiesis by an IdU pulse-chase experiment on the CyTOF carries with it significant qualitative advantages with respect to existing methods. We develop a combination of a pulse-chase IdU experiment and model-based data analysis to study hematopoiesis *in vivo* in wild-type adult mice. Our method allows to simultaneously track multiple populations, thereby giving a systemic view of differentiation, making full use of the power of mass cytometry. Our method can be applied on a variety of systems to generate systemic, quantitative comparisons, directly observing and quantifying differentiation.

Here we present our method, a combination of an *in vivo* mouse experiment and computational analysis and model fitting which yields direct observation of the differentiation flux during hematopoiesis. We first demonstrate the power of mass cytometry by identifying immune cell subsets by high dimensional clustering followed by plotting changes in their IdU profile over time. Then we illustrate our mathematical modelling approach by using conventional Boolean gating on bivariate scatterplots to track three different transitions in mouse bone marrow: neutrophil maturation, Pre-B to Immature-B cell development, and Immature-B to Mature-B cell development.

## Methods

### Experimental sketch

A schematic of the experimental plan is depicted in Fig. 1A. To track the differentiation of monocytes we injected IdU into the peritoneum of 48 C57BL-6J age-matched mice at 12 hour intervals. Mice are then sacrificed in cohorts of 16 on consecutive days, and their bone marrows were harvested, prepared as single-cell suspensions, stained with antibodies, and analyzed by CyTOF mass cytometry.

**Figure 1.**
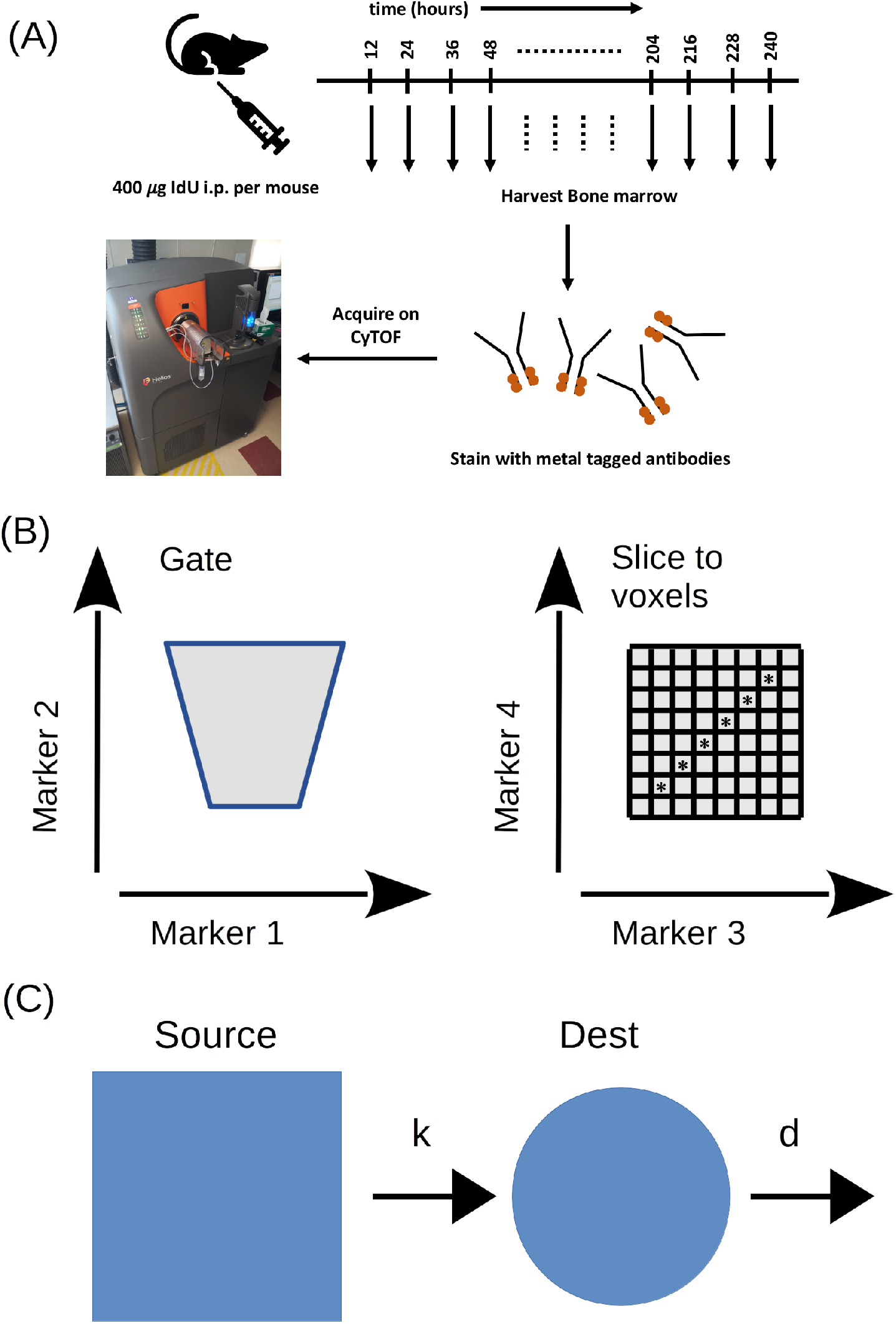
Summary experimental and modeling method: **(A)** Experimental setup: mice are injected *i.p*. with IdU, sacrificed at indicated time intervals for harvest of bone marrow, which is then stained with metal-conjugated antibodies followed by acquisition on a Helios mass cytometer. **(B)** For modeling differentiation along a phenotypic continuum, we zoom in on a transition of interest based on bivariate scatterplots and define a grid along its phenotype directions to create a map of voxels. We then focus on differentiation along the line defined by the first principle component (PC1) of the cell abundance distribution, marked by asterisks. **(C)** Schematic of pair-wise differentiation model and the transfer function equation which describes it.

### Mice

6-8 weeks old female C57BL/6J mice were used for the study. All animals received food and water *ad libitum*. All animal experiments were in accordance with the ethical approval of the institutional animal ethics review board.

### IdU Injection

IdU (*5-Iodo-2’-deoxyuridine*) was purchased from Sigma-Aldrich (ref. I7125). All mice were injected in their peritoneum with a solution of 0.4mg of IdU diluted in 200*μl* of sterile PBS, adjusted to a pH of 8.5-9. The injection times were set to sacrifice 16 mice per day over consecutive days, with each day covering as evenly as possible the time-range between 1 and 228 hours thus effectively representing a separate experiment. Each day included an un-injected control mouse in the cohort. Each time-point is represented by 2-3 different mice; the injection/collection regimen was structured to prevent batch effects in the pulse-chase dynamics.

### Cell preparation for CyTOF analysis

We euthanized the mice at various time points post IdU-injection, harvested and prepared their femoral bone marrow as single-cell suspension. Cells were then stained with metal-chelated antibodies according to manufacturer’s protocols (Fluidigm). Fc block antibody was included in the staining cocktail to minimize nonspecific antibody binding on Fc-expressing cells. The panel of antibodies can be found in the supplementary information.

### Data pre-processing

We used the data normalization package from the CyTOF manufacturer (Fluidigm) to correct for signal drift during acquisition. We then use manual gating to remove the calibration beads and dead cells (Cisplatin^+^) and to select singlet cells (peak at 2n for ^191^Ir-DNA intercalator). Further details of the gating strategy can be found in the supplementary information.

### High-dimensional analysis of mass cytometry data

To visualize the high dimensional data generated by CyTOF, we first project the data in a two-dimensional map by performing dimensionality reduction by t-Distributed Stochastic Neighbor Embedding (t-SNE) (26) using the median intensity values of all cell surface markers used in the experiment. The resulting t-SNE dimensions are then used to cluster the cells into leucocyte subpopulations by applying the spanning tree progression analysis of density normalized events (SPADE) algorithm to paint the clusters on the t-SNE map (35). Both t-SNE and SPADE were run in Cytobank (www.cytobank.org) using default parameters. The full IdU time course for all the obtained clusters are then plotted using Python.

### Defining a phenotypic continuum of differentiation

Classical FCM analysis involves displaying a density map of observed fluorescence variables two at a time, in a two-dimensional (bi-axial) plot, and then manually selecting sub-populations (gating) based on local abundance peaks. Such an approach is valid when dealing with large differentiation steps (19). More often than not, cells in fact differentiate and change their surface marker expression profile, in a more gradual and continuous manner.

Previously, our group dissected surface marker expression continua, gaining insight into the heterogeneous T-cell response to stimulus (20, 21, 22). For example, In Ref. (21), following FCM analysis, cells were binned according to their IL-2R*α* surface abundance in bins 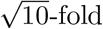 apart (i.e. 10^2^ – 10^2.5^, 10^2.5^ – 10^3^, etc.). The cells in each bin were analyzed for their pSTAT5 response to a range of IL2 doses, extracting a continuum of response sensitivities to IL2, which depends on IL2-R*α* levels. This method introduced the concept of *slicing* of the IL-2R*α* histogram to reveal a continuum of phenotypes. In the same spirit, here we employed a *slicing* method to study differentiation along phenotypic continua of leukocyte subpopulations within the bone marrow of mice. Here, however, instead of a one-dimensional grid, (dependence on a single surface marker), we divide a focus population to a two-dimensional grid of marker expression levels.

The two-dimensional slicing procedure is shown in Fig. 1B whereby each grid voxel would then be treated as a distinct population. Each voxel’s full time-series is then averaged over experimental replicates. Next, we define a one-dimensional direction in this phenotypic continuum by looking at the direction where the density distribution changes the most - namely, the first principle component (PC1) of the principal component analysis of the 2D density maps. We are now in a position to interrogate the differentiation dynamics along this path. For that purpose, we first develop our mathematical model and then fit it to the data.

### Mathematical model

Our IdU experimental setup presents a unique modeling opportunity to quantify the dynamics of differentiation and proliferation. Our modeling accounts for the simple transfer of the IdU from the proliferating populations to their progenies. Such transfer-function models are ubiquitous and their time-series representations have been well-studied in many scenarios (23). From the modeling perspective, our innovation lies in the application of this simple formalism to the question of cell differentiation, and the reliance on high-quality dynamics afforded by the CyTOF/IdU pulse chasing. In Fig. 1C, we show a schematic of the model which captures the flow of IdU from a “Source” to a “Destination” population. *k* is the rate going from the source population to the destination population; *d* is the rate cells leave our “Destination” population. The schematic in Fig. 1C implies an ordinary-differential-equation for the rate of change of IdU signal intensity,

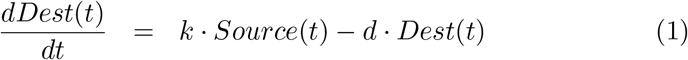

which when inverted produces the integral equation

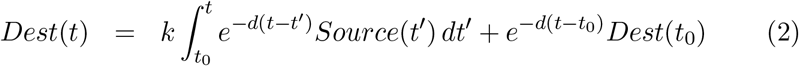

This integral equation takes the (experimental) source population’s IdU intensity, and fits values for *k* and *d* which best explain the destination’s IdU signal. To apply our model on the experimental data, we relied on the time-discrete version of the integral equation, where the integral is approximated as a sum of discrete time-points, a well-established technique (23). A similar mathematical structure was previously used to model genetic fate-mapping data (8, 24) but with crucial differences: (i) whereas the genetic fate-mapping captures the rise in label, experimentally it does not capture the subsequent decay that exists in IdU pulse-chase time-series. The direct observation of the decay thanks to the long-term monitoring of the IdU signal after a short pulse, better constrains the model. In Ref (13) it is precisely the arrest of the experiment before the decay can be measured in the Ly6C^low^ population that rendered those data less attractive to modeling; (ii) In Refs. (8, 24), homeostasis (steady-state) is assumed and is utilized to resolve the two rates *k, d* whose sum is the only experimentally-accessible observable (steady-state distribution of cells determines the ratio *k*/*d*). Conversely, our IdU pulse-chasing resolves both incorporation and decay, *k* and *d* are independent rates that are fitted independently, and the resulting values can be checked for compatibility with the homeostasis hypothesis. Hence, IdU pulse-chasing is advantageous as it affords stronger experimentally-constrained fit of our model.

### Data and code availability

All data used in this manuscript, including an MIFlowCyt checklist (25) is available in https://flowrepository.org/id/FR-FCM-ZYUV. All code used to process the data and generate the figures, including the fitting of the data to our model, is freely available in https://github.com/AmirErez/IdUPulseChase.

## Results

### Clustering of high-dimensional data allows for parallel visualization of differentiation dynamics of multiple immune cell types

In order to demonstrate the capability of mass cytometry to simultaneously track the differentiation dynamics of multiple immune cell types, we first projected the high-dimensional data into two-dimensional space using t-SNE as a dimensionality reduction tool (26). Subsequently, the cells in t-SNE space were clustered into distinct subpopulations by applying the SPADE algorithm (35). Fig. 2A shows a t-SNE map of 2-D projection of the obtained clusters based on the surface expression of individual markers. The clustering heatmap in figure 2B shows differential expression of cell surface receptors on the clusters that is used to identify classes of immune cells. In order to check for possible immune cell activation by IdU injection, we plot the median expression of surface MHC class II on a number of clusters over time. Figure 2C clearly demonstrates that the surface expression of MHC class II remains constant in different immune cell clusters over the entire time course of the experiment, thereby ruling out any potential change in differentiation dynamics as a consequence of cellular activation induced by IdU. Finally, we plotted the full time course of IdU positive cells for each of the identified clusters (figure 2D); these plots demonstrate that clusters are dynamically-diverse with peak in IdU uptake occuring earlier than 12hr (e.g. Cluster #11) or later than 72 hr (e.g. Cluster #22). Such analysis demonstrates the power of coupling IdU pulse chase with mass cytometry to observe differentiation dynamics of immune cells in the mouse bone marrow on a global scale.

**Figure 2.**
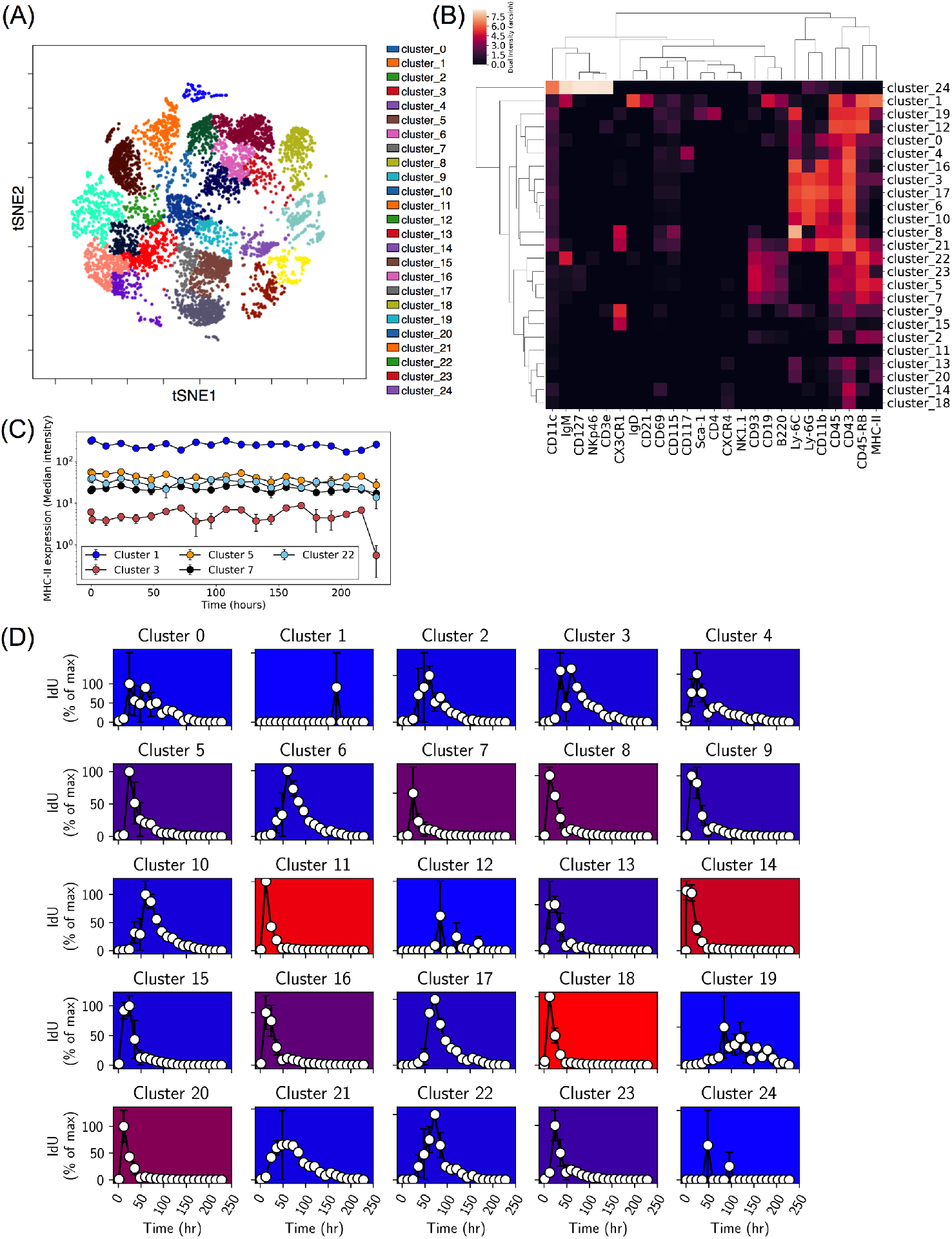
Global analysis of IdU pulse-chase by CyTOF. **(A)** t-SNE map of Live Single cells in the bone marrow demonstrating the clusters obtained by the SPADE algorithm. **(B)** Heatmap showing the expression patterns of cell surface markers used in the study across all the clusters obtained as in A. **(C)** MHC-II surface expression over time in select clusters demonstrating lack of immune cell activation by IdU injection. **(D)** Time dynamics of IdU uptake (normalized to maximum) for each leukocyte subpopulation. The color coding represents the range of IdU signal (red: high; blue: low).

### Phenotypic stability for biaxial gated leukocyte subpopulations

In what follows, we focus on particular populations of Neutrophils and B cells in the IdU time-series analysis. To further confirm a lack of any notable change in marker expression profiles across these studied populations, we examined the MHC-II levels for each mouse in the biaxial gating regime, as a function of time post IdU injection (Fig. 3). We found that, for 0.4mg of IdU as validated in our protocol, MHC-II expression does not vary with time in all leukocyte subpopulation. In particular, hand-gated neutrophils maintain lower MHC-II expression in comparison with the B-cells, as expected (Fig. 3). This strongly suggests that IdU injection in mice does not elicit any substantial inflammatory response. Additionally, for each population of interest, the markers used for studying differentiation dynamics were also found to be stable throughout the entire time series (Fig. 3).

**Figure 3.**
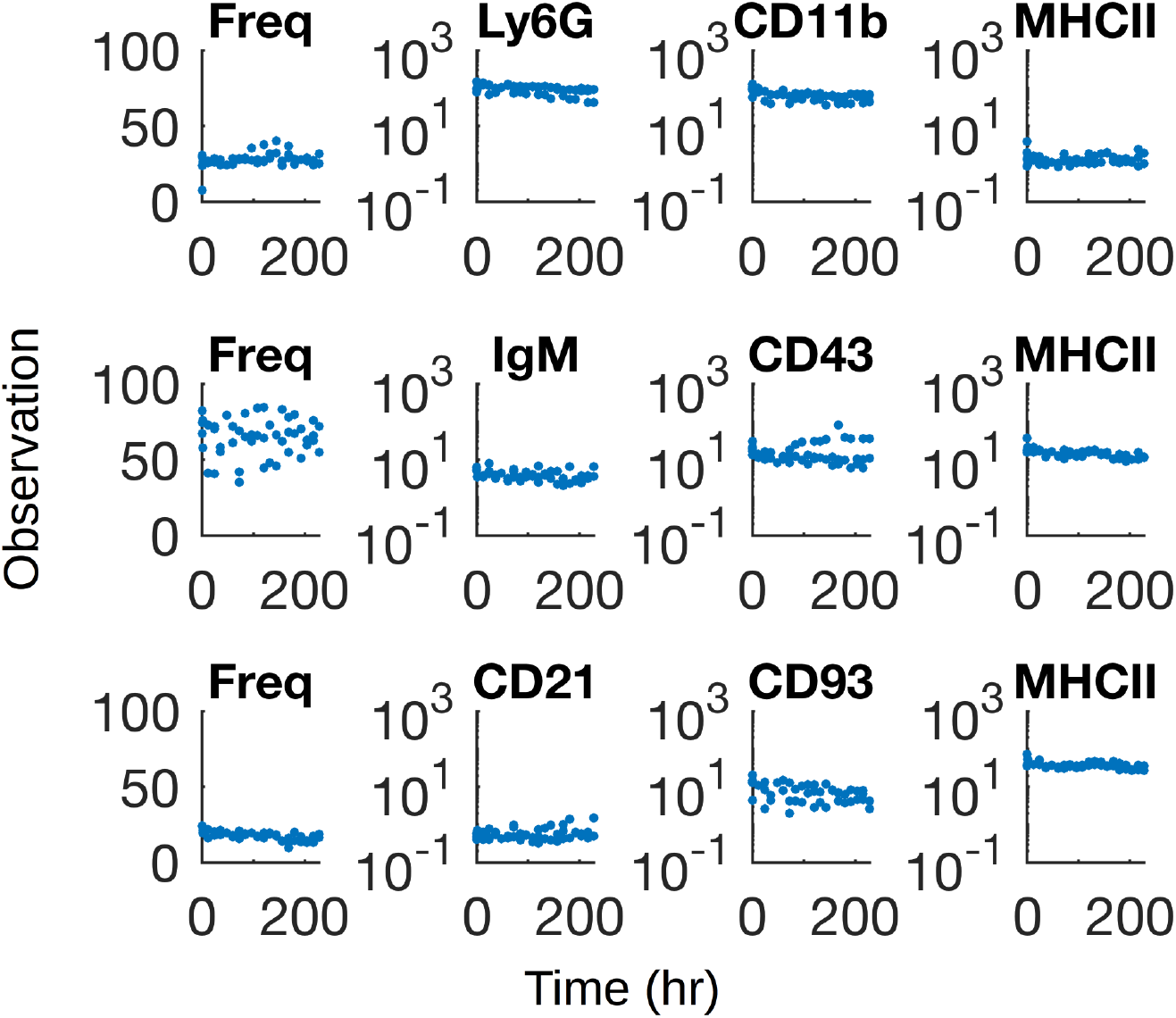
Phenotypic stability. Top row: neutrophils; middle: pre-B to immature B; bottom: immature-B to mature-B. None of the selected populations display any noticeable structure in their gate frequency (left column) nor any noticeable structure in their marker’s fluorescence intensity (two center columns) nor signs of inflammation in MHC-II expression (right column). This confirms that IdU injection does not perturb the bone marrow composition.

Taken together, these results support the temporal stability in leukocyte phenotype *i.e*. minimal inflammation throughout the IdU pulse-chase experiment. Thus, we demonstrate that the use of IdU pulse-chase allows one to monitor the dynamics of leukocyte differentiation while maintaining system homeostasis.

### Fitting the time-series on select groups of immune cells with known paths of differentiation

We then moved on to our primary objective, which was to illustrate the strength of our modeling methodology. To this end, we focused on individual tracks of differentiation within well-characterized cell lineages (e.g. Neutrophils, Immature B cells, and mature B cells), as defined by manual gating (see Supp Figures). For each lineage, we focus on a line in 2-dimensional marker space defined by the first *principal component* (PC1) of the cell count distribution (see Figs. 4B, 5B and 6B with the black asterisks). Each voxel (corresponding to a grid element as in Fig. 1B) is now treated as a separate subpopulation with its own time-series of IdU levels. We then decomposed the dynamics of differentiation into individual steps between nearest-neighbor voxels in the marker space. Meaning that, each voxel now has *progenitor* and *progeny* populations as the voxels above or below it along the line defined by PC1. We applied our modeling approach for the IdU dynamics within each subpopulation, using the pairwise fitting as applied to each subpopulation along PC1. In Figs. 4C, 5C and 6C, we compare the data (points) to the best fit according to the model (lines). The color code reflects the location along PC1 which each curve captures. Overall, our model captures well the dynamics of differentiation in the data, directly modeling and quantifying the dynamics of maturation within each cell lineage.

**Figure 4.**
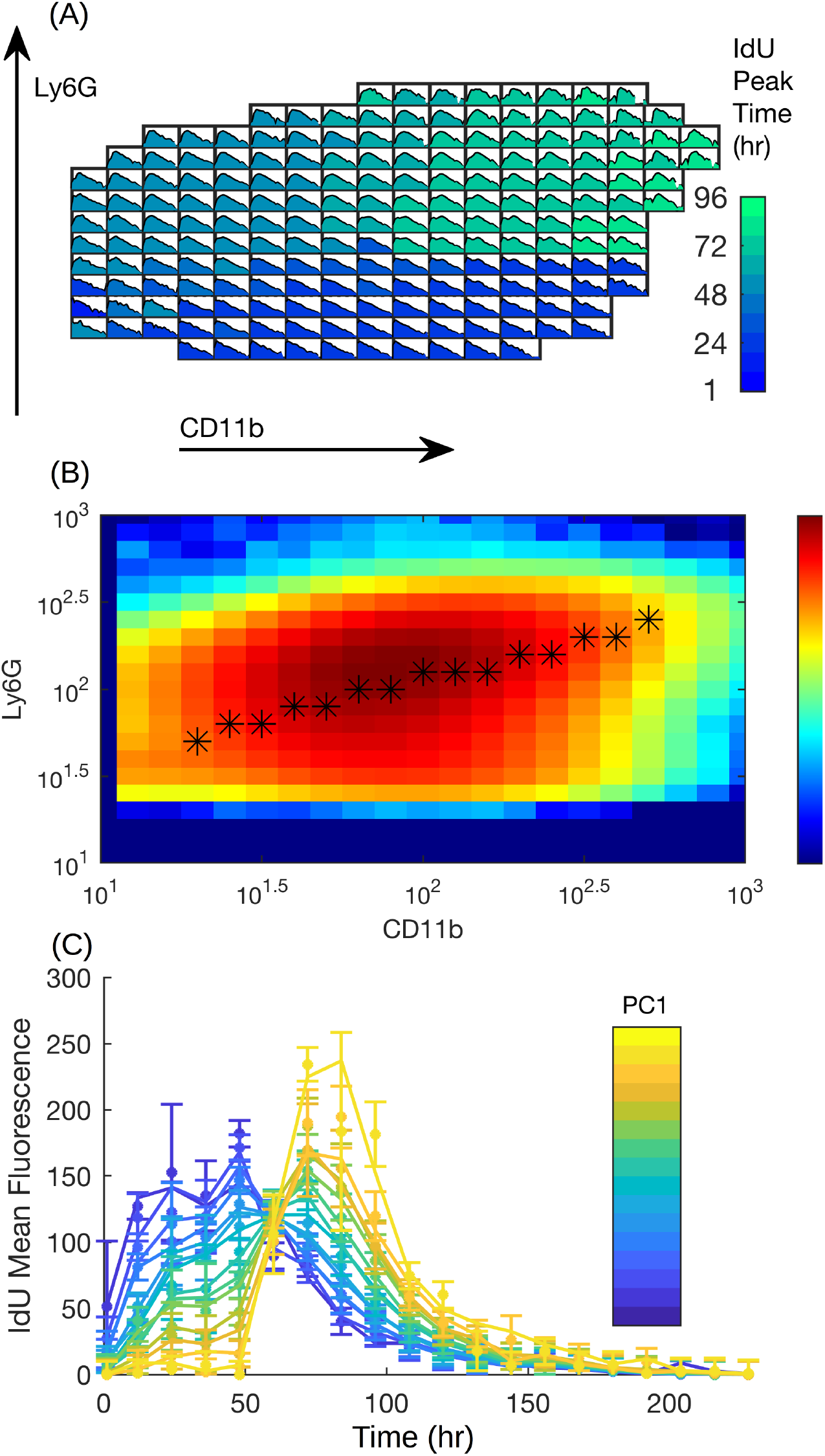
Neutrophil maturation. **(A)** IdU Time-series for each voxel, colored according to its peak IdU time. A voxel’s position corresponds to a range of Ly6G (vertical) and CD11b (horizontal) abundances which define the population. Each voxel shows an entire time-series (228 hours) of IdU mean-fluorescence intensity of the voxel’s population. **(B)** Density heatmap showing relative abundance for each voxel. The black asterisks mark the high-density differentiation path along the first principle component (PC1) of the density distribution. **(C) Time-series fit along PC1:** we follow the time-series for each of the voxels marked with an asterisk, showing both data (dots) and transfer-function (Eq. 1) fit (lines). Colors correspond to progression along PC1.

**Figure 5.**
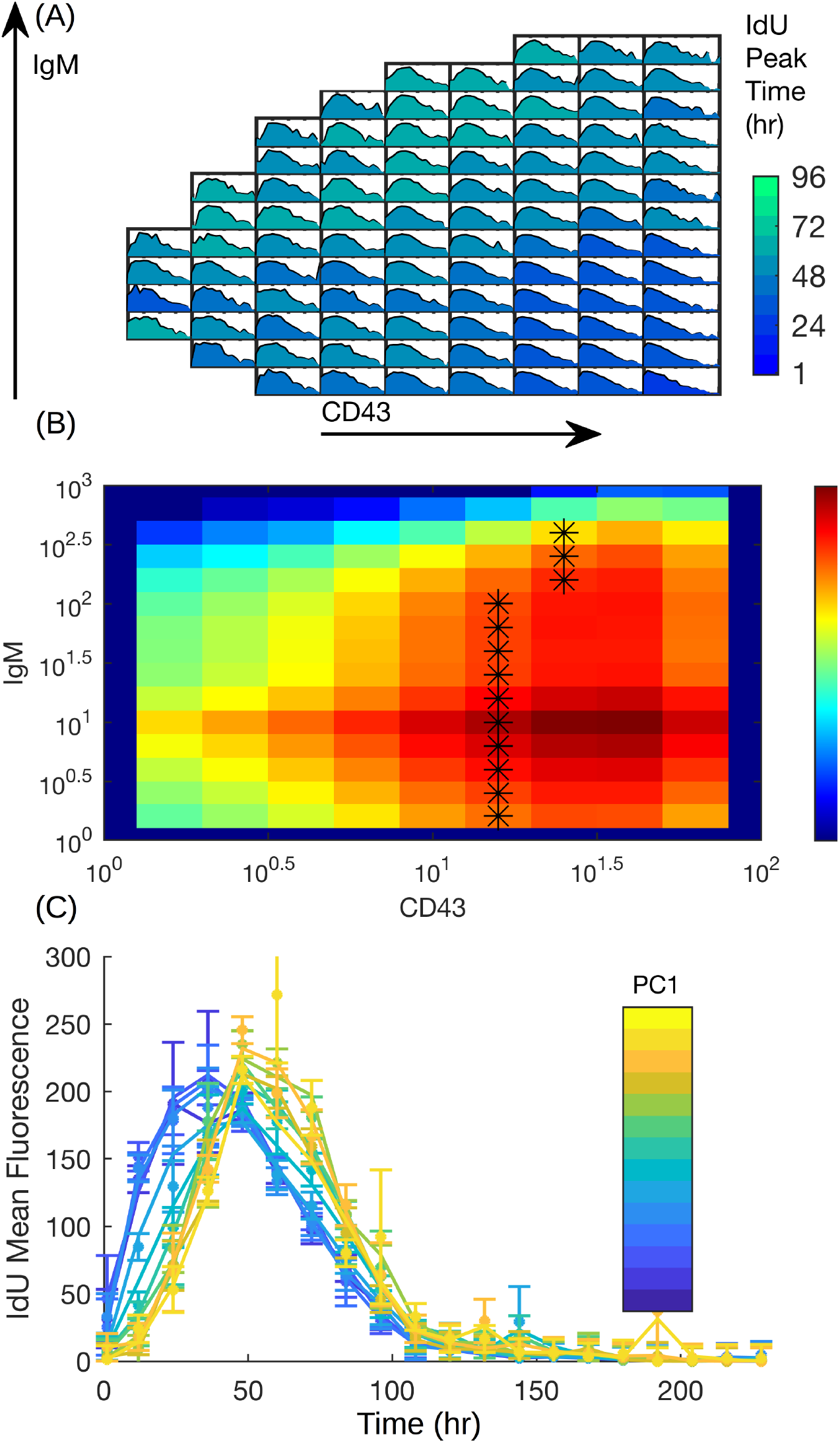
Pre-B To Immature B Cell transition. **(A)** IdU Time-series for each voxel, colored according to its peak IdU time. A voxel’s position corresponds to a range of IgM (vertical) and CD43 (horizontal) abundances which define the population. Each voxel shows an entire time-series (228 hours) of IdU mean-fluorescence intensity of the voxel’s population. **(B)** Density heatmap showing relative abundance for each voxel with asterisks showing PC1. **(C) Time-series fit:** data (dots) and transfer-function (Eq. 1) fit (lines). Colors correspond to progression along PC1.

**Figure 6.**
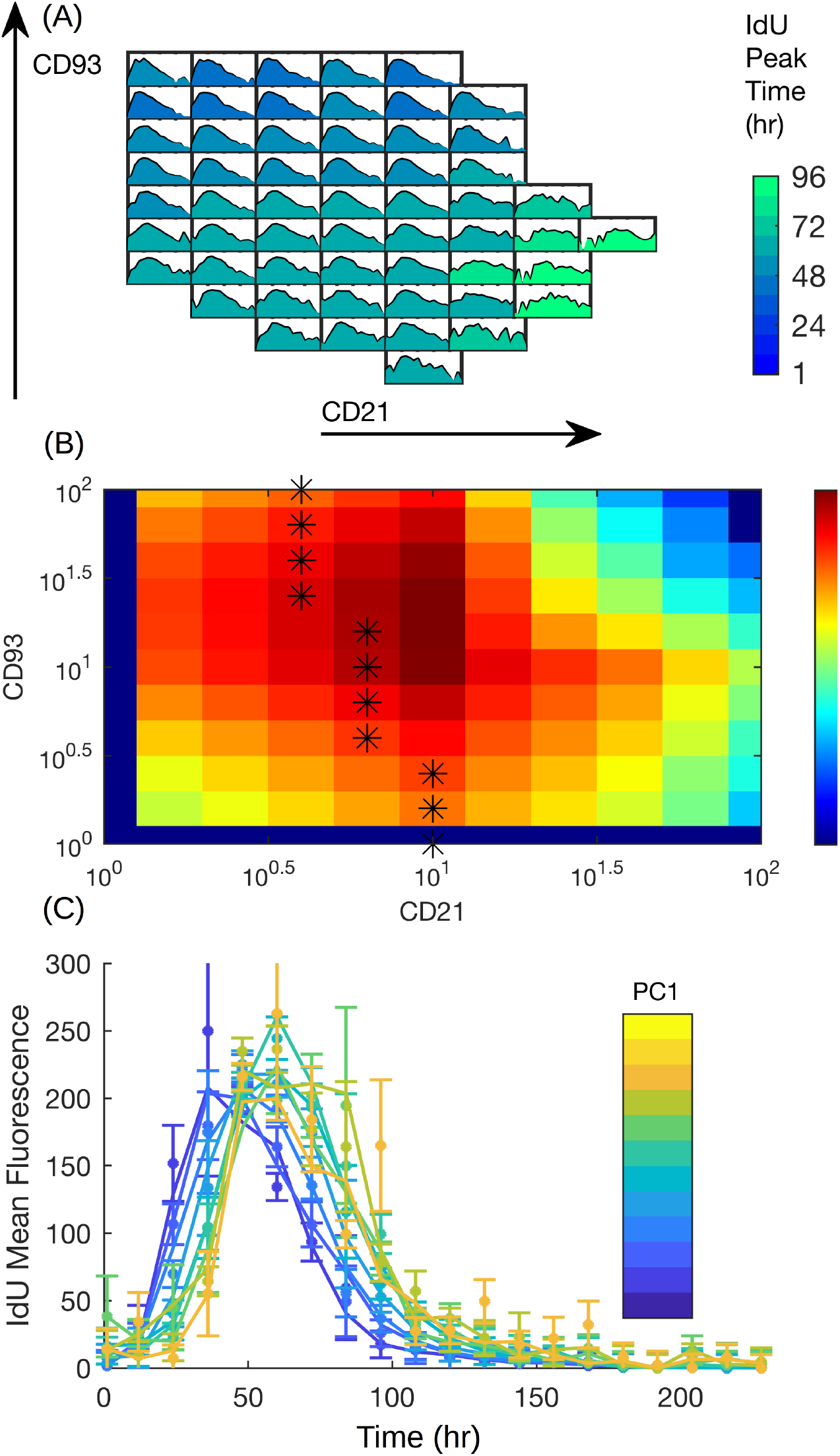
Immature to Mature B Cell development. **(A)** IdU Time-series for each voxel, colored according to its peak IdU time. A voxel’s position corresponds to a range of CD93 (vertical) and CD21 (horizontal) abundances which define the population. Each voxel shows an entire time-series (228 hours) of IdU mean-fluorescence intensity of the voxel’s population. **(B)** Density heatmap showing relative abundance for each voxel with asterisks showing PC1. **(C) Time-series fit:** data (dots) and transfer-function (Eq. 1) fit (lines). Colors correspond to progression along PC1.

#### (i) Neutrophils

We begin by observing the maturation dynamics of neutrophils. As demonstrated in Fig. 4A, the IdU peak time increases as we progress diagonally across from Ly-6G^lo^/CD11b^lo^ to Ly-6G^hi^/CD11b^hi^ populations, suggesting an increase in both Ly-6G and CD11b surface expression as neutrophils mature, as reported previously (27). Moreover, proliferating neutrophils were found to express lower Ly-6G than non–proliferating ones, as indicated by the noticeably decreased Ly-6G expression amongst IdU time points (Fig. 4A). This observation is also in accordance with a recently published study (18), where the authors reported decreased Ly-6G surface expression on proliferating neutrophils. Modeling the time-series by following the trajectory along the first principal component axis of the density distribution reveals two major peaks of differentiating populations. As can be seen in Fig. 4B, Ly-6G^lo^/CD11b^lo^ neutrophils peak at around 40 to 50 hours, whereas the Ly-6G^hi^/CD11b^hi^ populations peak at approximately 80 to 90 hours. The peak times estimate the time differentiation took since the main contribution from proliferating progenitors. Thus, it takes approximately 40 hours for the neutrophils to upregulate CD11b and Ly-6G as they mature along PC1, as marked by the asterisks in Fig. 4B.

#### (ii) B cells

The dynamics of B cell development in mice is a well-studied question (28, 29, 30). We took advantage of this to further validate the efficiency of our approach and to recapitulate known biology. CD43 is a marker that is expressed on all B cell precursors - Pre-Pro-B, Pro-B, and Pre-B cells (31, 32, 33). As Pre-B cells differentiate into immature B cells they gradually gain surface IgM (33). Fig. 5 shows the transition dynamics of Pre-B cells to immature B cells, which displays two distinct differentiating cell types - an early CD43^pos^/IgM^neg^ Pre-B cell population followed later by IgM^pos^ Immature B cells Fig. 5C. Interestingly, the two differentiation peaks are separated by only a few hours, suggesting a rapid transition from precursors to immature B cells.

We then sought to model the transition of immature B cells to mature B cells (Fig. 6). Immature B cells are high on CD93 and sIgM while low on CD21 (28). As they mature, they progressively downregulate expression of CD93 while upregulating expression of CD21 (28). Here again, such an elementary step in differentiation can be delineated using a one-dimensional path in the (CD21, CD93) plane along which we can monitor the dynamics of IdU pulse-chasing. As depicted in Fig. 6C, the immature B cells appear to peak in IdU uptake around 50 hours post-IdU pulsing, followed by a decline. As observed with the pre-B to immature B cell transition, the mature B cell differentiation peak almost overlapped with the immature B cells, indicating a very rapid transition.

It is by no means obvious that Eq. 1 would indeed fit the time-series data so well. From that, we can conclude that the differentiation process convincingly follows along PC1 and that it does not involve significant nonlinear effects, e.g., secretion of iodine by the cells. This serves as a powerful proof of concept for the IdU pulse-chase framework we propose here. Moreover, we are now in the position to compare the estimated values of the various *k* and *d* parameters. In Fig. 7A we plot *k* and *d* for the three compartments (circles) together with the *k* = *d* line (dashed). It is apparent that *k* ≈ *d* with *d* ≥ *k*. The latter is not surprising: by definition our model allows *k* from only the progenitor (“Source”) to go to the progeny (“Destination”), whereas *d* is not limited in such a way. Specifically, *d*, which quantifies the rate cells exit a subpopulation, includes differentiation, as well as other processes, e.g. cell death, resulting in *d* ≥ *k*. Indeed, it is this flexibility, allowing *k* and *d* to be independent, that differs from a related modeling approach (8). Previously, Busch *et al*. (8) imposed a strict steady-state relation between *k* and *d* since, unable to observe the decay of the signal, they cannot estimate *d* independently. Of further note is that the rates appear comparable across the three transitions, suggesting that it is not a particular cell type transition that is fast or slow. Rather, for each cell type, *slicing* it to subpopulations reveals a range of differentiation rates within that broadly defined compartment.

**Figure 7.**
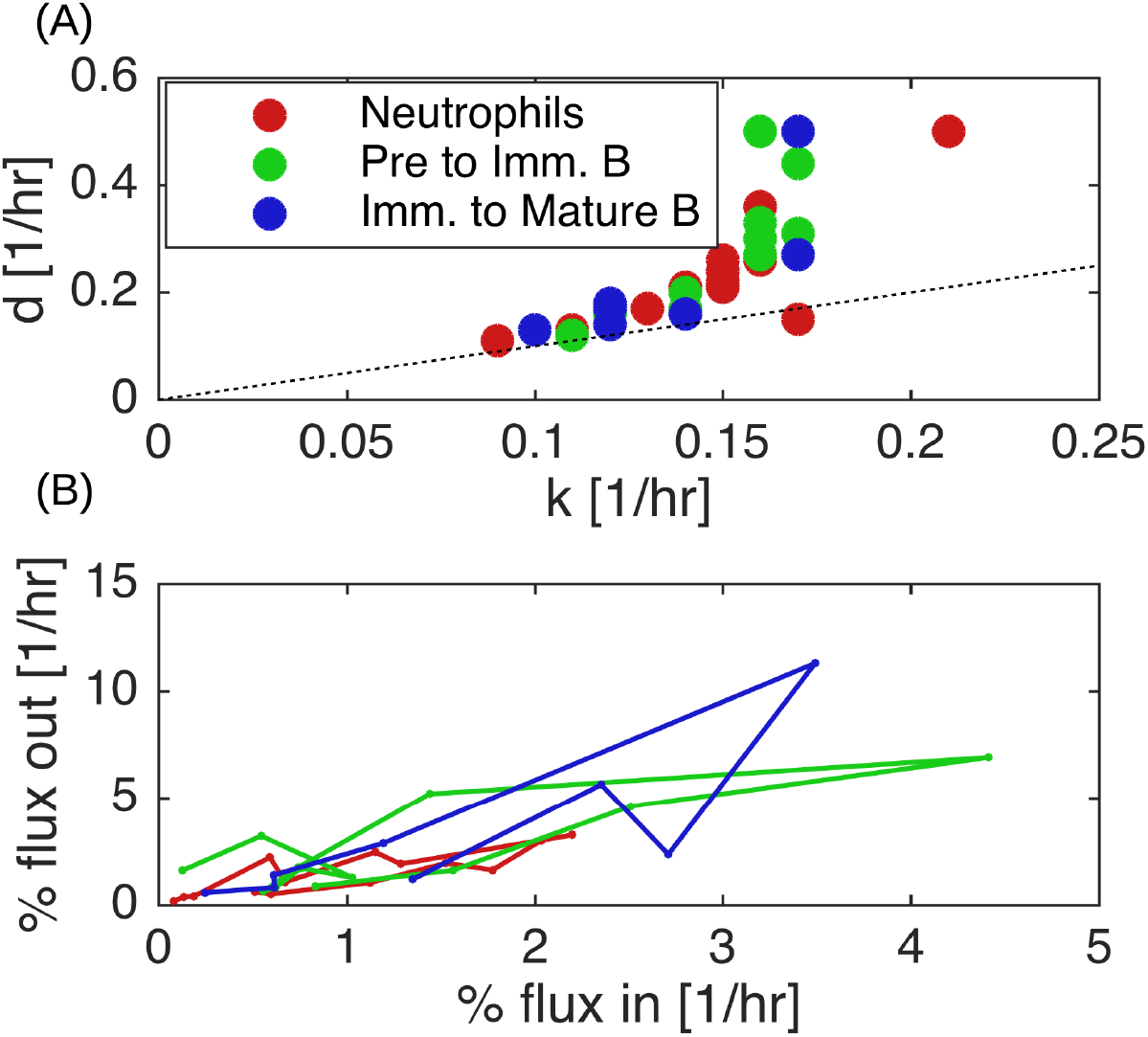
Comparison of differentiation rates. **(A)** Bare rates *k, d* for the neutrophil maturation (red), pre-B to immature B (green), and immature B to mature (blue). Dashed black line corresponds to *k* = *d*. **(B)** Differentiation fluxes along PC1. % Flux in (out) is calculated as k (d) multiplied by the cell count fraction along the progenitor (progeny) of the respective voxels.

The modeling strategy presented here can also be used to glean information about the cellular fluxes. The rates *k, d* shown in Fig. 7 quantify how *fast* a cell transitions between neighboring voxels, and are agnostic to *number* of cells in question. In contrast, the flux measures the *fraction of cells per hour* that transition. To estimate the fluxes, we multiply *k* by the fraction of cells that belong to each progenitor and *d* by the progeny’s fraction (normalized to the total cell count along PC1). We plot the fluxes in Fig. 7B, with the lines connecting the trajectory along PC1. As is apparent, for subpopulations at the edge of the compartment (e.g. the lowest Ly-6G^lo^ and CD11b^lo^ cells in Fig. 4B), the differentiation flux is low, consistent with their smaller abundance fraction. In the three transitions in this manuscript, namely the neutrophils and the two B cell transitions, we note a pattern of maximal flux around the center of the transition, with the B cells replacing a larger overall percentage per hour than the neutrophils.

Together, the above results clearly demonstrate that our method, which combines the power of mass cytometry with mathematical modeling can faithfully recapitulate previously reported facets of dynamics of neutrophil and B cell differentiation in the mouse bone marrow (27, 28, 29, 30, 33). For example, we demonstrated that immature Ly-6G^lo^/CD11b^lo^ neutrophils peaked in their IdU uptake earlier than mature Ly-6G^hi^/CD11b^hi^ neutrophils, hence confirming the natural dynamics of differentiation in the neutrophil compartment. Similarly, our analysis of B cell development revealed a clear progression of pre-B cells to mature B cells via immature B cells by sequential loss and simultaneous gain of key cell surface markers that are known to be associated with B cell development (28, 29, 30, 33).

## Discussion

Discrete gating by Boolean logic is standard practice for identifying cell populations of interest among heterogeneous cell samples by FCM. However, expression of a certain marker on a given cell type is highly variable even within a given population defined by that marker. To better capture this cell-to-cell heterogeneity we *sliced* cell populations into a grid of subpopulations, with each subpopulation a voxel on a bivariate plot. This allowed us to model relative changes in marker expressions as the cells transition from one marker state to another.

We introduced here a methodology combining IdU pulse-chasing, CyTOF monitoring and mathematical modeling to experimentally monitor the dynamics of differentiation. Using this approach, we quantitatively monitored the dynamics of three major cell type transitions during hematopoiesis in the bone marrow of adult mice. This enabled us to measure the real dynamics of differentiation rather than a pseudo-time of purely-phenotypic approaches.

We confirmed the simultaneous increase of Ly-6G and CD11b expression in the neutrophil compartment as the cells mature. Our observations are in agreement with a previously published report (34). In addition, we observed two temporally well separated peaks of differentiating populations in the neutrophil compartment with the peak differentiation of fully mature neutrophils happening at around day 4.

In contrast, our analysis suggested that the maturation of B cells from their precursors happens on a much shorter time scale with very little time lag between development of early and late stages. Our methodology will be of particular interest when studying how such differences in maturation dynamics between immune cell types shape an immune response, e.g. during an inflammatory response.

Mass cytometry is fast gaining popularity for immune cell phenotype analysis because of the number of simultaneously measurable markers it allows without running into the technical difficulty introduced by fluorescence spillover between channels in classical fluorescence-based cytometry. In the present study, we took advantage of our monitoring of *in vivo* IdU uptake, and our direct detection of iodine by mass cytometry, to study hematopoietic dynamics in mouse bone marrow at a more global scale. This enjoys the advantage of easy preparation and minimal perturbation of the sample by reagents required to access the nucleus in its FCM counterparts, EdU and BrdU. Although studying progression of immune cells through stages of cell cycle by IdU incorporation followed by analysis by mass cytometry has been introduced before for *in vitro* setting (17), its use to study differentiation *in vivo* has been limited. In a recent study, Evrard *et al*. used *in vivo* administration of IdU to phenotypically and functionally distinguish proliferating and non-proliferating populations of immune cells (18). Here, we built on these two papers to perform IdU pulse-chase in live animals over multiple time points and to address the dynamics of differentiation.

To analyze multiple time-points, we introduced a simple mathematical modeling scheme. Our model benefits from the fact that tracking a tracer molecule (here IdU) is not a biological phenomenon (with all implied complexity), but rather, is a simple physical phenomenon readily captured by a set of linear pairwise transfer relations. And so, it is the combination of, which provides the strength and flexibility for monitoring hematopoietic differentiation: the IdU allows CyTOF acquisition with its technical advantages; The slicing objectively treats surface protein expression heterogeneity without imposing on it a binary classification; The time-series and its model-based analysis provide the direct quantification of the differentiation dynamics. The simplicity of each of these three aspects is what, when taken together, renders them a general and effective technique. Having established our method, one possible extension of the work presented here will be to automate the manual gating steps. Naturally, such automatic gating would need sufficient discourse and proper validation, which is why, for simplicity, we used manual gating in this study. Another interesting extension of our method would be to investigate differentiation trajectories beyond the first principle component (PC1). Although PC1 is a natural choice, since by definition is captures the dominant direction in surface marker heterogeneity, it is by no means the only choice. Indeed, one might model differentiation beyond a one-dimensional trajectory, modeling instead the multi-dimensional flow of cells through phenotype space, though such models will require a significant theoretical effort.

To conclude, we introduced a quantitative framework for analyzing the dynamics of cell maturation that can be leveraged to study differentiation of multiple immune cell types in parallel. It will be interesting to see whether such a broad and detailed mapping of hematopoiesis at homeostasis or under inflammation would reveal unexpected systemic differences in mouse strains and/or well-used transgenic models.

## Acknowledgments

This work was supported by Human Frontier Science Program grant LT000123/2014 (Amir Erez), the Gordon Moore foundation and by the Intramural Research Program of the NCI, NIH. We thank April Huang for her help with tagging the mice. We thank Brian Sellers for expert advice with mass cytometry. The CyTOF Facility at the Center for Human Immunology is supported by NIH intramural funding (AI001226-01).

