## Supplemental Information for "Quantifying the dynamics of hematopoiesis by *in vivo* IdU pulse-chase, mass cytometry and mathematical modeling"

### Supplementary methods

#### Gating strategy

To focus on well-known leukocyte differentiation steps and to illustrate the strength of our method, we focused on transitions of interest by usual bi-axial gates. We defined a lineage-negative population as $Sca1^{-}CD11b^{-}CD11c^{-}CX3CR1^{-}CD3e^{-}NK1.1^{-}CD4^{-}CD117^{-}$. Then, we define three sub-populations as follows: (i) $Lineage^{-}Ly6G{+}CD19^{-}$ neutrophils ; (ii) $Lineage^{-}CD11b^{-}Ly6G^{-}CD19^{+}IgD^{-}CD43^{-}$ Pre and Immature B-cells; (iii) $Lineage^{-}CD11b^{-}Ly6G^{-}CD19^{+}IgM^{+}IgD^{lo}$ Immature and Transitional B-cells.

In Fig. 1 we see the bone-marrow gating strategy; In Fig. 2 we see the lineage- gating for the B-cells.

#### List of Antibodies

All antibodies were purchased from Fluidigm and used at dilution ration 1:100, apart from Cisplatin and DNA which were used at 1:1000.

| Antigen | Metal | Clone | Cat | Lot |
|---|---|---|---|---|
| CD45 | 89Y | 30 F11 | 3089005B | 2471519 |
| Ly-6G | 141Pr | 1A8 | 3141008B | 931506 |
| CD11c | 142Nd | N418 | 3142003B | 2751404 |
| CD69 | 143Nd | H1.2F3 | 3143004B | 2301412 |
| CD115 | 144Nd | AFS98 | 3144012B | 2821412 |
| CD45-RB | 145Nd | C363.16A | 3145012B | 2311507 |
| CD43 | 146Nd | S11 | 3146009B | 2691401 |
| CD19 | 149Sm | 6D5 | 3149002B | 301503 |
| IgD | 150Nd | 11-26c.2a | 3150011B | 31409 |
| IgM | 151Eu | RMM-1 | 3151006B | 3001411 |
| NKp46 | 153Eu | 29A1.4 | 3153006B | 2211306 |
| CD11b | 154Sm | M1/70 | 3154006B | 1531405 |
| CD93 | 158Gd | AA4.1 | 3158015B | 2321405 |
| CXCR4 | 159Tb | L276F12 | 3159030B | 2881606 |
| B220 | 160Gd | RA3-6B2 | 3160012B | 331513 |
| Ly-6C | 162Dy | HK1.4 | 3162014B | 1041608 |
| CX3CR1 | 164Dy | SA011F11 | 3164023B | 1201614 |
| CD3e | 165Ho | 145-2C11 | 3165020B | 3181410 |
| CD21 | 168Er | 7G6 | 3168010B | 2371409 |
| Ly-6A/E | 169Tm | D7 | 3169015B | 331514 |
| NK1.1 | 170Er | PK136 | 3170002B | 901512 |
| CD4 | 172Yb | RM4 5 | 3172003B | 2791505 |
| CD117 | 173Yb | 2B8 | 3173004B | 331524 |
| MHC-II | 174Yb | M5/114.15.2 | 3174003B | 791514 |
| CD127 | 175Yb | A7R34 | 3175006B | 141524 |
| Cisplatin | 195Pt | N/A | 201064 | 1801605 |
| DNA | 191/193Ir | N/A | 201192A | 1331616B |

#### Supplementary figures

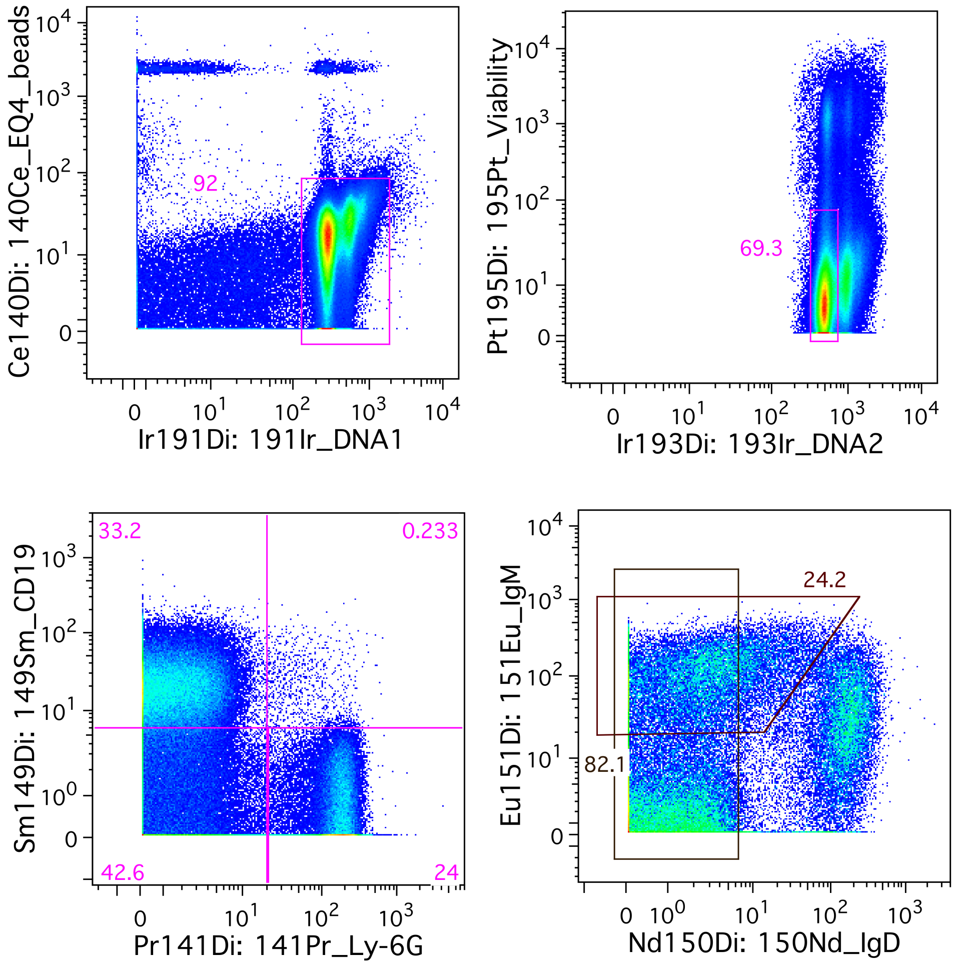


**Figure 1. Gating the bone marrow:** **(A)** Selecting cells as Beads-DNA+ then **(B)** live single cells by Cisplatin-DNA gating, followed by **(C)** Neutrophils defined as Ly6G+CD19- and B-cells as Ly6G-CD19+. The B-cells are further selected for lineage- according to Fig. 2 and then divided to **(D)** (i) Pre to Immature transition ($IgD^{-}/IgM^{-/+}$ gate) and (ii) Immature to Transitional ($IgM^{+}/IgD^{-}$ gate).

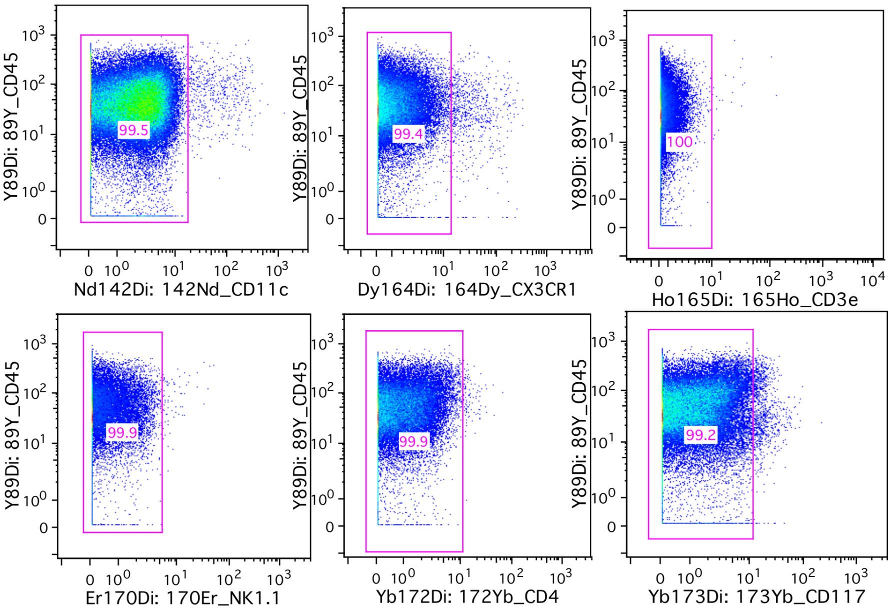


**Figure 2.** Lineage- gates: B-cells are selected for Ly6G-CD19+lineage- according to Fig. 1 and the lineage gates above.
